# Expansion Sequencing of RNA Barcoded Neurons in the Mammalian Brain: Progress and Implications for Molecularly Annotated Connectomics

**DOI:** 10.1101/2022.07.31.502046

**Authors:** Daniel R. Goodwin, Alex Vaughan, Daniel Leible, Shahar Alon, Gilbert L. Henry, Anne Cheng, Xiaoyin Chen, Ruihan Zhang, Andrew G. Xue, Asmamaw T. Wassie, Anubhav Sinha, Yosuke Bando, Atsushi Kajita, Adam H. Marblestone, Anthony M. Zador, Edward S. Boyden, George M. Church, Richie E. Kohman

## Abstract

Mapping and molecularly annotating mammalian neural circuits is challenging due to the inability to uniquely label cells while also resolving subcellular features such as synaptic proteins or fine cellular processes. We argue that an ideal technology for connectomics would have the following characteristics: the capacity for robust *distance-independent labeling, synaptic resolution, molecular interrogation, and scalable computational methods*. The recent development of high-diversity cellular barcoding with RNA has provided a way to overcome the labeling limitations associated with spectral dyes, however performing all-optical circuit mapping has not been demonstrated because no method exists to image barcodes throughout cells at synaptic-resolution. Here we show ExBarSeq, an integrated method combining in situ sequencing of RNA barcodes, immunostaining, and Expansion Microscopy coupled with an end-to-end software pipeline that automatically extracts barcode identities from large imaging datasets without data processing bottlenecks. As a proof of concept, we applied ExBarSeq to thick tissue sections from mice virally infected with MAPseq viral vectors and demonstrated the extraction of 50 barcoded cells in the visual cortex as well as cell morphologies uncovered via immunostaining. The current work demonstrates high resolution multiplexing of exogenous barcodes and endogenous synaptic proteins and outlines a roadmap for molecularly annotated connectomics at a brain-wide scale.

## Introduction

The availability of high-resolution brain connectivity maps will dramatically enhance our ability to understand how neural activity drives function and behavior, as well as accelerate the development of the next generation of therapeutics. Light microscopy (LM) is ideal for mapping brain circuits because of its ability to rapidly image thick tissue samples, to integrate biomolecules as varied as proteins, nucleic acids, and small molecules, and to distinguish these species using a wide multispectral bandwidth (Osten and Margrie 2013). In order to map connectivity, the molecular information must enable labeling and recovery of individual synapses, linking those synapses to the correct pre- and post-synaptic cells, and scalable recovery of many cells across large volumes. A central challenge to this approach is the encoding of information: using traditional fluorescent probes such as antibodies or GFP, there are simply not enough labels to uniquely number the large number of cells that may overlap in a given volume, even when used in clever combinations (Cai et al. 2013; Veling et al. 2019).

An appealing solution to this problem is to label cells with unique identifiers composed of nucleic acid sequences, instead of spectral fluorophores. This approach maximizes the number of potential labels because it provides exponential diversity; a set of nucleic acid sequences of length N provides for 4^N^ unique labels, and even a short 20-nucleotide sized pool of oligos can trivially label over one trillion unique cells (exceeding the number of neurons in the mouse or human brain). Our labs have explored several methods for barcoding cells in this manner over the past decade, including use of programmable zinc fingers (Mali et al. 2013), self-editing DNA barcodes using CRISPR-Cas9 (Kalhor et al. 2018) and directly labeling neurons with unique RNA barcodes that are trafficked to synaptic sites (Kebschull et al. 2016).

In the MAPseq technique, each labeled neuron is nominally infected with a single RNA barcode, which is expressed at high copy number in the soma and also transported to distal projections using a protein carrier. Across a population of labeled neurons, the long-range projections of each neuron can be identified by dissecting potential targets and sequencing the RNA barcodes transported to each region (Han et al. 2018; Kebschull et al. 2016). This method enables massively parallel readout of long-range projections, and can be integrated with in-situ sequencing to enable direct readout of both cell body locations and endogenous gene expression (BARISTAseq; X. Chen et al. 2018; Sun et al. 2021). A limitation of these techniques is their spatial resolution: diffraction-limited LM lacks the precision necessary for reliable identification of individual synapses. However, recent advances in integrating in-situ nucleotide imaging with resolution improvements from direct expansion of the target tissue provide a potential path towards multimodal molecular connectomics using scalable LM techniques (ExSeq; Alon et al. 2021).

Together, techniques like BARISTAseq and ExSeq provide a possible foundation for a new class of experiments that use combinatorial labeling and super-resolution microscopy to map neuroanatomical connectivity at a precision and scale previously impossible. Here we present ExBarSeq, which combines multiplexed barcode labeling, in-situ sequencing, sequential immunostaining, Expansion Microscopy (ExM), and novel computational analysis to provide a comprehensive platform for mapping neuronal connectivity. In this approach, we first use barcoded Sindbis virus libraries to uniquely label a population of neurons with RNA barcodes, which are expressed in both the soma and in distal projection or putative synaptic sites. Second, we recover these barcodes with high spatial resolution using an improved protocol for combining ExM and in-situ sequencing, resulting in clear labeling of important morphological features. In combination with the experimental procedures, novel algorithms are employed to computationally demix ambiguous or overlapping barcodes, resulting in high-resolution localization of anatomical markers for each cell. We show that this approach successfully identifies the location of cortical neurons in labeled brains, sparsely labels a subset of the overall cellular morphology, and labels sites adjacent to specific synapses. We conclude that this method – if supplemented by improved barcode expression from the Sindbis virus or other method of genetic delivery – is potentially sufficient to create comprehensive maps of neuronal connectivity at the brain-wide scale.

## Results

### Experimental Protocol

RNA barcodes virally delivered to cells in vivo through Sindbis viral libraries using the MAPseq technique have been shown to provide diverse RNA labeling of thousands of cells at nearly clonal multiplicity of infection (**Figure 1A**). Individual virions carry 30-nt RNA barcodes, which are amplified within a target cell after infection, and an expressible GFP for morphological tracing. The diversity of these nucleic acid labels provides digital cellular labeling, up to 4^30^ different barcodes. To increase labeling density, the barcode is highly expressed as a subgenomic mRNA, with subcellular transport enabled by a protein carrier. In this experiment, mouse visual cortex was injected with MAPseq viral libraries (Kebschull et al. 2016), and tissues were subsequently prepared using standard protocols compatible with a broad range of LM techniques.

**Figure 1.**
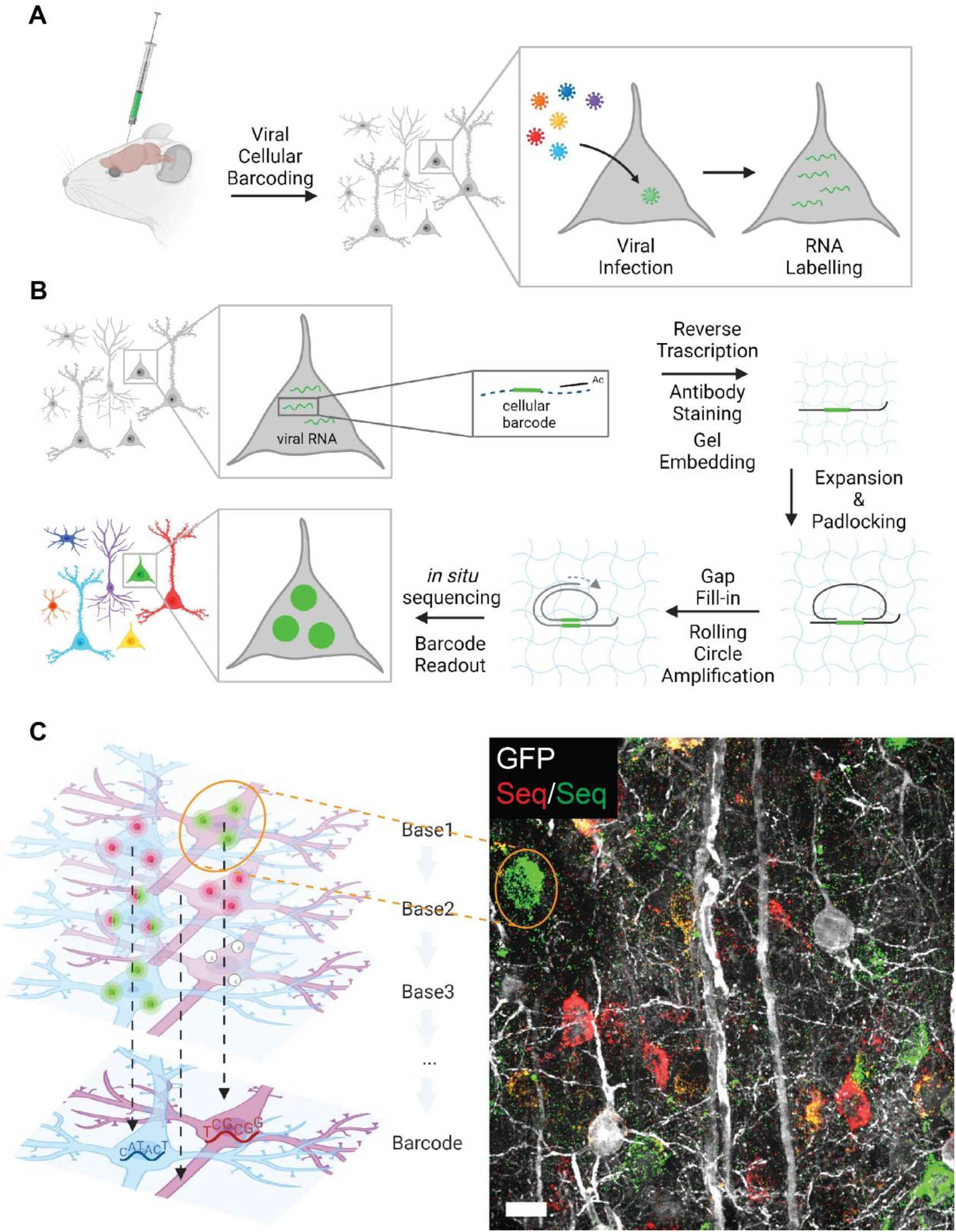
Experimental pipeline of ExBarSeq. (A) Sindbis vector libraries are injected into an adult mouse brain where each vector expresses many copies of one distinct RNA barcode per neuron (B) The ExBarSeq library preparation protocol involved converting RNA barcodes into cDNA which, along with antibody stains, get anchored into an expandable hydrogel. Amplification of cDNA using rolling circle amplification creates amplicons that can be subjected to in situ sequencing. (C) In situ sequencing processes base-by-base in 3D, for 6 rounds. The image on the right shows representative data for an individual sequencing round. Scale bar is 50 *µ*m post-expansion.

To recover the identity and spatial location of these RNA barcodes, we used a variant of ExSeq chemistry (Alon et al. 2021) that captures each RNA barcode and embeds it in a hydrogel for subsequent in-situ sequencing. The RNA barcode is recovered through reverse transcription (RT) of subgenomic mRNA, using a primer that is modified with both locked nucleic acid (LNA) nucleotides to maximize binding affinity to a conserved region of the barcode and a 5’ acrydite group to enable radical polymerization for stable capture within the ExSeq hydrogel matrix (**Figure 1B**). The result is cDNA containing the barcode that is anchored into the hydrogel and ready for subsequent amplification. We reasoned similar chemistry could be used for concurrent antibody labeling.

To integrate RNA barcodes with morphological information and synaptic protein labels, tissues were then immunostained with antibodies targeted to the synaptic proteins Bassoon and Homer1, as well as to the virally-introduced GFP. Each antibody was conjugated to an oligonucleotide tag, which serve two purposes: first, they carry polymerizable acrydite end groups that embed into the hydrogel scaffold; second, they enable reversible imaging of each probe through multiplexed in situ hybridization (ISH). Following this labeling step, cellular proteins as well as protein antibodies were removed via proteolysis, which provides for robust optical clearing. Expansion of the hydrogel scaffold is due to a combination of electrostatic repulsion forces within the polymer network and osmotic pressure gradients while utilizing anchoring molecules to maintain the spatial relationship between embedded probes. This expansion is then “frozen” at 3.3x by re-embedding the sample in a neutral hydrogel and chemically modifying the remaining acrylate functional groups to remove their intrinsic negative charge which allows for enzymatic activity on the spatially anchored molecules (Alon et al. 2021).

Neuronal barcodes, present in both soma and distal processes, are recovered using gap-filling padlock probes and amplified using rolling-circle amplification into ∼250 nm cDNA nanoballs to enable in-situ sequencing-by-synthesis (SBS) with high SNR (X. Chen et al. 2018). In addition to SBS, antibody targets are labeled via nucleic acids that can be amplified and imaged using target-specific FISH probes. The FISH probe for the barcode amplicons (cDNA nanoballs) doubled as a sequencing primer because the dye was placed on the 5’ end leaving the 3’ end available to prime the sequencing steps, enabling two-color Illumina Sequencing chemistry and multi-round 3D registration for intact tissue.

We first performed in-situ sequencing of the expanded sample to interrogate neuronal barcodes in the context of cellular morphology via antibody-labeled GFP (**Figure 1C**). For each two-color SBS round of ExBarSeq imaging, we acquired two microscope channels of in situ sequencing, one channel for amplicon localization, and one channel for GFP-labeling of Sindbis-infected neurons. As both the antibody and neuronal barcodes are labeled with molecular tags (i.e. the sequence of the padlock probe and the sequence of the FISH probe, respectively), we were able to recover and co-localize both modalities in a single sequencing operation. More generally, this approach combines recovery of two libraries – one set of designed barcodes used for combinatorial ISH labeling and one set of random barcodes used for massively multiplexed labeling of neurons. This protocol enables consistent capture of all targets though tissues up to a thickness of 30 µm (pre-expansion). Imaging of slices up to 100µm pre-expansion thickness resulted in robust barcode labeling but incomplete staining from antibodies, therefore all data shown here were conducted on 30*µ*m thick slices.

RNA barcode nucleotides could be clearly determined for soma-associated amplicons and allowed for labeling of hundreds of neurons within intact cortical tissue (**Figure 2A**). MAPseq cortical injections produced a high expression bolus with GFP trafficking visible through six layers of cortex (**Figure 2B, 3B**). Imaging a single neuron at 40x magnification, we observed barcode amplicons inside dendritic spines (**Figure 2C**). Furthermore, we observed a general pattern in which barcode density decreased farther away from the cell body, with amplicon density higher in perisomatic spines than in distal spines.

**Figure 2.**
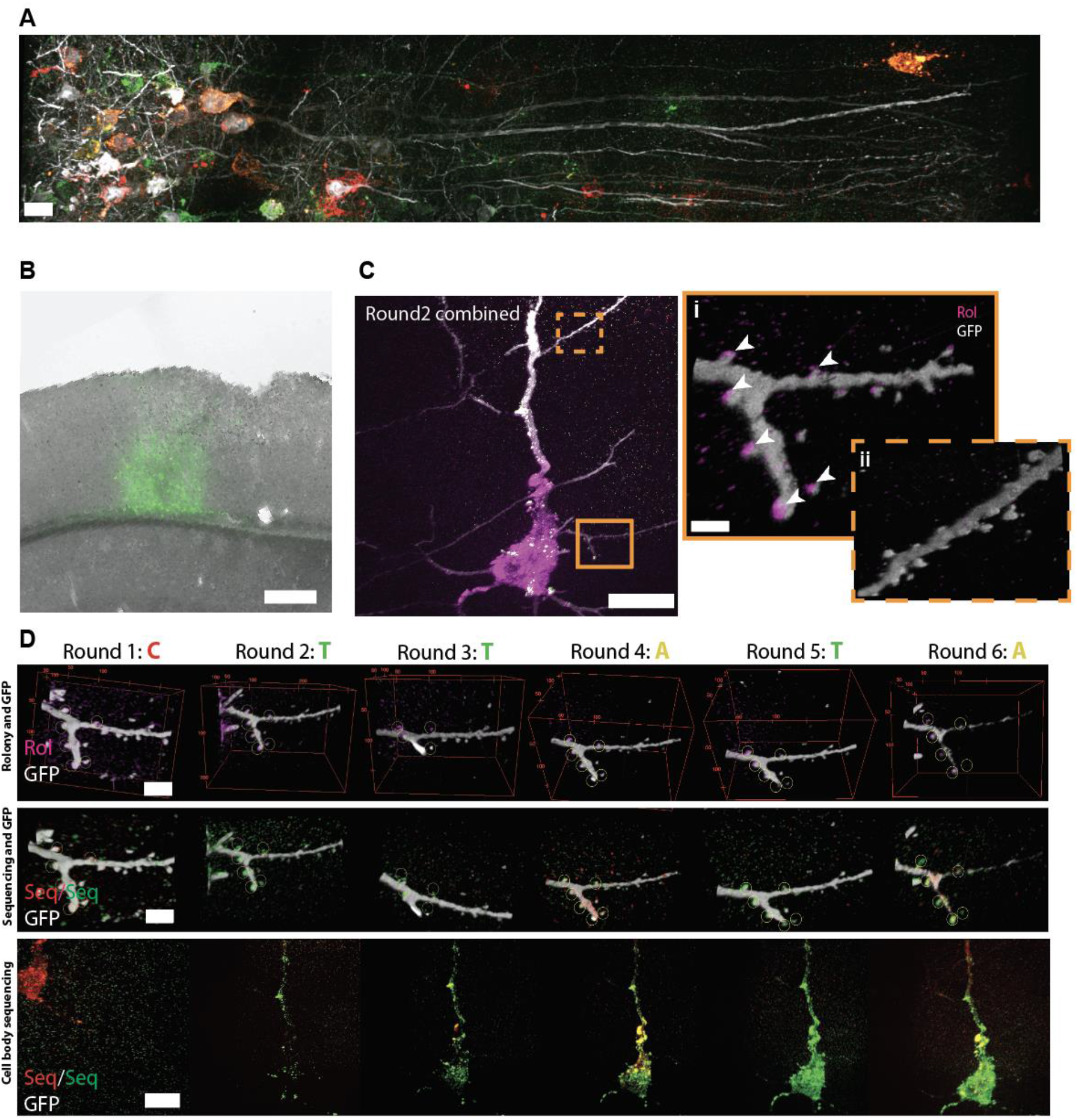
ExBarSeq resolves RNA barcodes inside dendritic spines. (A) Representative image of a round of sequencing within the injection bolus throughout multiple layers of mouse cortex, dense labeling is observed in the cell bodies. (B) Widefield image of Sindbis library injection site in the mouse cortex. GFP fluorescence is overlaid with a brightfield image. (C) Viewing a single barcoded neuron for one round of in situ sequencing with a 40x objective demonstrates clear morphology of dendritic spines as well as rolony barcode amplicons (Rol) located adjacent to synapses (i, arrowheads). Distal dendritic spines did not have observable barcode amplicons (ii). (D) Visualizing the same neuron as in (C) across six rounds of in situ sequencing in both the dendritic section (Ci) and the cell body. Yellow circles around dendritic spines correspond to the same locations as arrowheads in (Ci). Note these images were not registered, so imaging Round 1 is offset by about 50 *µ*m. Scale bars are 20 *µ*m (A), 250 *µ*m in (B), and 20 *µ*m for the cell body image in (C) and (D), then 2 *µ*m for the dendrite view in (Ci, Cii and first two rows of D), post-expansion except for (B) which was not expanded.

As a practical note, overall duration of the ExBarSeq experimental protocol is determined by the amount of imaging that was performed and is otherwise insensitive to the physical volume of the sample. For this experiment, large tiling was performed at 20x magnification to reduce imaging time. Each brain sample can be subjected to multiple rounds of sequencing, enabling reliable identification of subcellular barcodes: for example, we show six rounds of sequencing of barcode CTTATA, clearly visible in both the cell body and in peri-somatic dendritic spines (**Figure 2D**).

### Computational Pipeline

Imaging of intact 3D cortical samples using ExBarSeq can generate large multi-dimensional datasets that are challenging to manage using existing pipelines. In order to interrogate an entire cortical column within the injection site we imaged 14 fields of view at 20x magnification using 4 channels to a depth of 100*µ*m post-expansion (total imaging volume of the expanded gel 1.5mm x 0.4mm x 0.1mm, in 4 channels), then imaged that same volume 6 times for 6 rounds of sequencing (**Supplemental Figure 1)**. The total dataset amounts to 12TB of imaging data, a data size challenging to render, register, and reconstruct (**Figure 3A**).

**Figure 3.**
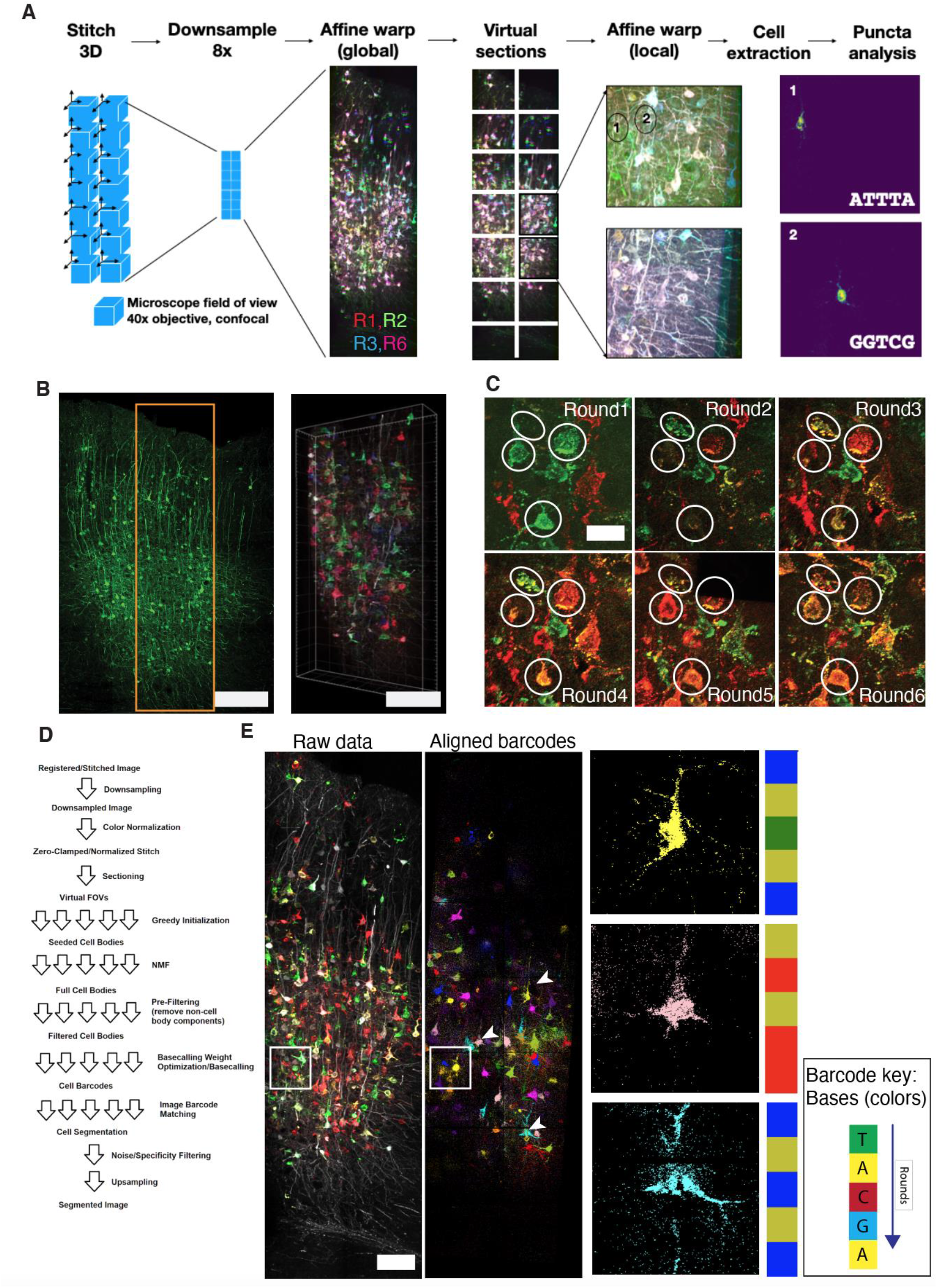
The ExBarSeq image processing pipeline integrates data across fields of view and sequencing rounds to identify distinct cellular barcodes. (A) Each sequencing round is stitched in 3D before being downsampled and registered with a sample-wide affine transform: Sequencing rounds 1, 2, 3 and 6 are shown in different colors and overlaid: white indicates overlapping signal of well-registered parts of the sample. Virtual fields of view (vFOV) are then created from the full, roughly aligned original data and then each vFOV is registered again at full resolution, from which the cellular barcodes can be automatically extracted from the cell bodies. Two sample extracted cells are shown. Pre-expansion confocal image of the injection site (left) and the post expansion 3D image of the volume that was in situ sequenced (right) (C) Representative data across six in situ imaging rounds - for clarity, only the two sequencing channels (red and green) are shown. (D) Conceptual diagram of the NMF-based barcode extraction method to create the dictionary of barcodes present in the data (E) Results of the NMF based barcode discovery and alignment across the cubic millimeter dataset. White arrow denotes the position of the cells shown at higher resolution in (B), the four circled cells in A are visible as three yellow cells and a light blue cell in the pseudocolored barcode image. Scale bars are 250 *µ*m in (B), 20*µ*m post expansion in (C) and 100*µ*m post expansion in (E).

One common challenge for *multiple field of view* (FOV) imaging is the alignment of distinct FOVs in 3D to create a contiguous image that reflects the underlying physical sample. Additionally, the challenge is increased for *multiple round imaging* methods such as in situ sequencing, because the 3D volume of the sample deforms slightly across imaging rounds (for instance, due to forces arising from temperature, osmolarity, ionic strength, and shear stress arising from solute flow). For this study, registration was particularly challenging because of the 1000x scale-difference between the required spatial resolution for synaptic imaging (∼100nm) and the spatial extent of the sample (∼1mm) (**Figure 3B**), and because physical deformation of a freely floating hydrogel is non-linear rather than affine.

To resolve the ExBarSeq challenge for *multiple round, multiple FOV* imaging experiments, we implemented a scalable registration approach that utilizes two affine transforms: for each round of sequencing, each FOV was first locally aligned to its neighboring FOVs using TeraStitcher (Bria and Iannello 2012) to create a complete image volume that is not registered to other rounds. This stitched image volume was then downsampled in TeraStitcher and processed to derive an affine approximation using the ExSeqProcessing library (Alon et al. 2021) of the entire sample’s deformation relative to the first round of imaging. This sample-wide approximate registration then allowed us to create “virtual” FOVs, or vFOVs, which were subsections of the stitched image and small enough for memory-efficient processing at full resolution using ExSeqProcessing.

While *multi-round, multi-FOV* has been done before (Murray et al. 2015), the previous solution utilized a non-linear warp to be calculated across the entire sample that required the entire image be loaded into memory, which is infeasible for larger image volumes. Without generating vFOVs, the 2TB stitched composite image from one sequencing round (13090 × 3892 × 400 pixels) would have been too big for full-resolution image processing. Our vFOV approach has the benefit for easy parallelization and utilizes the single-voxel accuracy of the ExSeqProcessing registration system. The resulting registered images allowed us to easily track cells across sequencing rounds, confirming the data was ready for more complex analysis. For the sample discussed in this paper, we analyzed 5 of the 6 sequencing rounds due to one round having a sample handling error (**Supplemental Figure 1**). The result from this registration is that every pixel inside the image references the same point in biological space across six sequencing rounds, allowing for RNA barcode analysis per barcode amplicon (**Figure 3C**).

In order to map the morphology of each neuron, we sought to sequence all RNA amplicons in the sample, not just the amplicons in the cell body where the expression was strongest. To do so, we first built a dictionary of barcodes present in cell bodies. This step is necessary because, unlike the barcodes in FISH-labeled antibody probes, neuronal barcodes are randomly chosen from a high-diversity library and are not known in advance. To extract these barcodes de novo, we developed a pipeline based on non-negative matrix factorization (NMF) of vFOVs (**Figure 3D**) that required neither hyperparameter turning nor previous training data (**SI Methods**). Inspired by successful NMF-based pipelines (S. Chen et al. 2021, 2022) for fluorescent calcium reporters (Pnevmatikakis et al. 2016; Giovannucci et al. 2019), this algorithm iteratively extracted cell bodies that expressed barcodes of a minimal complexity across sequencing rounds, then converted sequential NMF components into 2-color Illumina encoding back into the ACTG base using a discriminative loss function (De Brabandere, Neven, and Van Gool 2017). Because of the relatively short length of the sequenced barcodes (5 of 30 bases), we enforced a uniqueness filter that automatically removed any barcodes that were expressed in multiple neurons within the imaged sample, and also rejected barcodes with somas outside the image volume. In the example slice analyzed here, this approach recovered a library of 63 barcodes, as detected by somatic expression, that were then analyzed on a per-amplicon basis for potential synaptic contacts.

To map amplicons from distal projection sites onto the library, we then matched the color-space signals of the full set of amplicon voxels identified using the ExSeqProcessing toolkit, excluding barcodes with low complexity barcodes (eg, AAAAA). The result is 50 distinctly labeled neurons with visible cell bodies, which are visualized in pseudocolor in **Figure 3E**. Importantly, this approach allowed us to locate and sequence barcodes that were trafficked outside of the soma into neuronal processes, while linking those barcodes to identified soma in the sample. Thus, a relatively simple unsupervised computation method on ExBarSeq data is enough to enable automated identification and localization of barcodes throughout the sample. Representative barcoded neuronal cell bodies discovered by the computational pipeline are shown in **Figure 3E** and the complete set of 50 extracted neurons is shown in **Supplemental Fig 2**.

### Analysis

We next asked whether identified barcodes could serve as fiducial markers for reconstruction of proximal neuronal morphology across the intact sample (**Figure 4A**). Although limited by the density of barcode expression from current-generation Sindbis construct, which does not completely fill distal projections, we explored whether relatively sparse labeling could enable recovery of proximal morphology for individual neurons, as well as aggregate distributions for more distal projections. Amplicons in the neuropil rich area of Layer 2/3 and showed spatial patterns corresponding to neuronal somas and local projections (**Figure 4B**), but the signal was insufficient for guiding the recovery of complete neuronal morphology.

**Figure 4.**
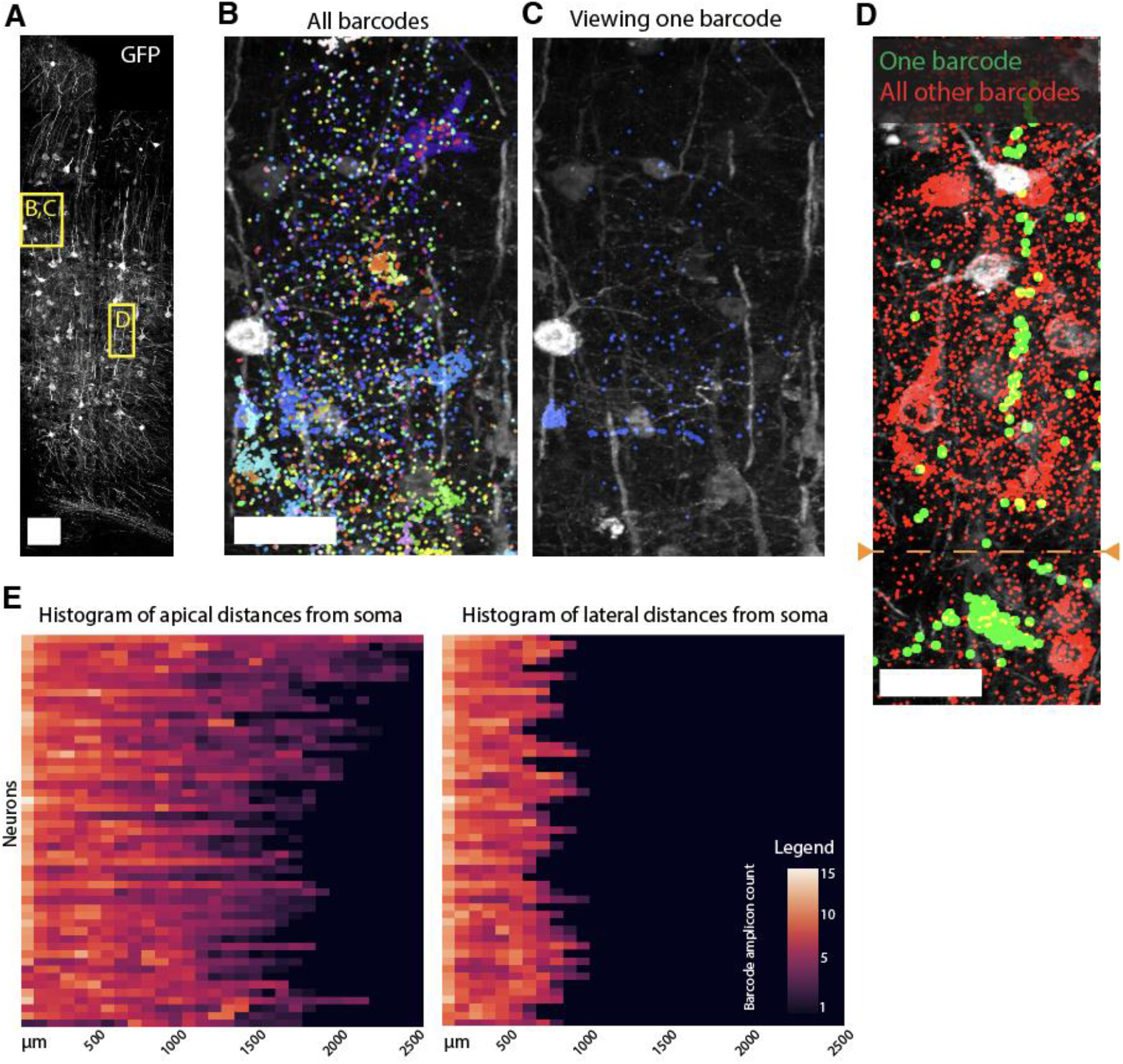
Automatically identified barcodes can be studied in situ for distal trafficking into neuropil. (A) GFP expression in the cortical sample illustrates the location of studied barcode trafficking locations. (B) Extracted barcodes from the neurons in the neuropil-rich Layer 2/3 indicate high barcode expression in cell bodies but lower barcode presence in the neuronal processes. Each dot is a barcode amplicon colored by barcode identity (C) Visualizing just one barcode’s spatial pattern indicates potential distal trafficking and, by lack of a GFP signal, illustrates the the barcode’s sub-cellular anti-correlation with GFP. (D) An RNA barcode signature indicates a clear spatial pattern indicating the presence of a neuron. One neuronal barcode, shown in green with its soma at the bottom of the image, demonstrates the expected apical projection in the cortex and can be visualized concurrently with all other aligned barcodes shown in red. The orange dotted “FOV boundary” denotes the point at which two adjacent FOVs were stitched successfully. (E) The statistical patterns of apical and lateral spatial distribution of RNA barcodes relative to the centroid of the Soma matches expected spatial distribution of predominantly apical projections. Scale of 100 *µ*m in (A) and 25 *µ*m in (B,C,D).

It is worth considering what improvements would be required to enable EM-level reconstruction in an ExM experimental context. First, ground-truth reconstruction will require co-localization with a morphological signal, GFP in this case. Here, this morphology recovery is currently limited by the known anticorrelation between barcode mRNA and GFP expression in virally labeled neurons (**Figure 4C**), which is likely driven by the suppression of translation by Sindbis proteins nsP2/nsP3 (Gorchakov, Frolova, and Frolov 2005). Still, we observe neuronal barcodes expressing the expected apical projection morphology up to ∼200 um distance from the Soma **(Figure 4D**). In parallel with the work shown here, there have been additional developments on both computational and experimental fronts to support ExM-based connectomic reconstruction (M’Saad et al. 2022; Yoon et al. 2017).

For more distal projections, a quantitative analysis of recovered barcodes confirms several expectations of the predominantly pyramidal-cell morphology which we observed in the somas **(Figure 3E)**: (1) typical morphologies show strong expression in a compartment corresponding to the apical dendrite of a pyramidal neuron, spanning over 700*µ*m (pre-expansion); (2) an overall alignment of barcodes favors the apical/basal alignment of pyramidal neurons by a factor of ∼3x (**Figure 4E**) (Spruston 2008). More comprehensive reconstructions will be enabled by improvement in the Sindbis virus, both to increase barcode count and resolution of the known anticorrelation in expression of the barcode and GFP.

### Synaptic Localization

Recovering synaptic labels is critical to mapping neuronal connectivity, but the small size of synapses renders them unresolvable using conventional, diffraction-limited imaging. Our use of Expansion Microscopy allowed direct resolution of these features. In this experiment, a subset of synapses were identified by immunolabeling with Bassoon and Homer1 as pre- and post-synaptic markers, and visualized using in-situ sequencing of ISH probes in expanded tissue (F. Chen, Tillberg, and Boyden 2015). Although these proteins do not generalize to all synapses, they serve as useful demarcations of neural junctions.

Using the ExBarSeq protocol, we visualized barcodes in conjunction with both endogenous synapse staining and GFP labeling to visualize cell morphologies (**Figure 5A**). We observed barcodes in close proximity to synaptic markers within labeled cells, with barcodes observed mostly near postsynaptic sites as determined by the directionality of the synaptic markers, but also observed at presynaptic sites (**Figure 5B**). This result supports the possibility of obtaining a full connectivity graph using ExBarSeq, limited currently by the challenge of improving the density of barcode expression. Still, the technological combination of antibody stains, highly multiplexed exogenous barcodes and ExM may be a path to connectomic interrogation of a mouse brain at 100x the speed of electron microscopy (EM) based connectomics (**Figure 5C**).

**Figure 5.**
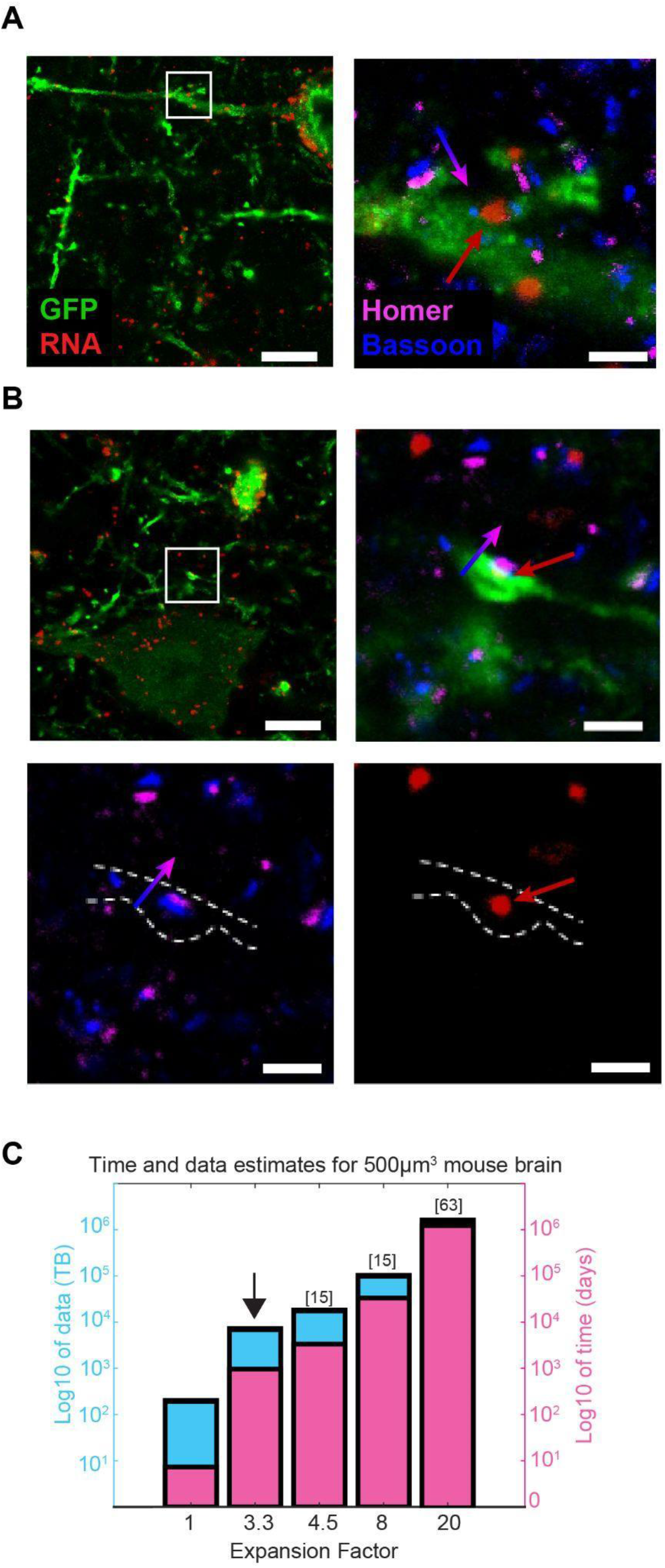
ExBarSeq visualizes barcodes colocalized with antibody stains to endogenous synaptic markers. (A) Examples of clearly distinct synapses as detected in 3.3x expanded ExSeq gel with a library preparation of RNA barcodes (amplicons in red) co-stained with homer (magenta), bassoon (blue) and GFP (green) and imaged on a 40x laser-scanning confocal. The barcode amplicon (arrowhead) in the dendrites is in close proximity to a synapse. (B) Images of a barcode amplicon at an axon terminal. Separate images of only-synapse and only-barcodes included with morphology annotation for clarity. Scale Bars: 20 *µ*m, 5 *µ*m post-expansion (zoom in). Red arrows show the location of RNA barcodes and blue/magenta arrows show the directionality of the synapses Time and data estimates for optical imaging of an intact mouse brain with a 40x objective across multiple expansion factors. Arrow indicates the expansion factor used in this study, which corresponds to 7.2 petabytes and 2.65 years of continuous imaging per round of molecular multiplexing. While other expansion factors have been demonstrated, 3.3x presents a balance of clearly distinctive synapses while being conceivably practical with sufficient microscopes.

## Discussion

The present study represents a step forward in using cellular barcoding for the all-optical readout of neural circuits in mammalian brain tissue. Because of the dense cellular architecture of the brain and the branched and polarized morphology of neurons, many unique channels are needed to be able to distinguish cell projections that are overlapping and in close proximity. We reasoned that by labeling cells with RNA as opposed to fluorophores, labeling diversity can rival or exceed the number of cells in the mammalian brain. In this regime, every cell would get its own barcode and therefore all cells will be identifiable within the brain tissue. If such RNA barcode labeling were combined with synaptic annotation and morphological stains, brain connectomes could be rapidly acquired using light microscopy.

The minimal connectome is a data structure that ascribes every synapse to its pre- and post-synaptic neurons. For a well designed brain mapping technology, capturing additional endogenous information would be a minimal marginal cost in addition to the morphological segmentation challenge, and such biological information would be essential for developing new understandings of circuit-level neurobiology. It is known that synapses are molecularly diverse with receptor types giving rise to large differences in physiology (O’Rourke et al. 2012; Hafner et al. 2019; Zhu et al. 2018). Therefore, molecular annotations of endogenous molecules to elucidate subcellular states should be considered an essential component for the future of connectomics.

We argue that an ideal all-optical technology for connectomics would have the following characteristics: robust *distance-independent labeling* (cellular barcoding), *synaptic resolution, molecular interrogation, and simple, self-contained computational analysis* (ie, no reliance on training data) (Table 1). Distance-independent labeling is important so that neurite segments can be identified and attributed to a cell body even if it is not possible to continuously trace the entire neuron. Especially in consideration of the hundreds or thousands of brain slices that may be necessary toward a single connectome, some amount of sample handling errors are bound to occur and so the technology must be robust in application. Synaptic resolution is important to see connections and also to attribute molecular properties to synapse-level morphological features. Finally, we wish for the technology to be usable inexpensively and equally applicable for both for future large-scale centralized mapping facilities and individual labs with targeted neurobiological inquiries.

**Table 1:** Summary of critical connectomic technology attributes and their status to date. We identify four critical characteristics for future optical brain mapping technologies. This demonstration succeeds at three of the characteristics but is unable to accomplish distance-independent labeling.

| Critical attribute | Description | Demonstrated here | Future work |
| --- | --- | --- | --- |
| Distance independent labeling | Every neuron must have a molecular barcode that links all components of interest back to its soma, independent of distance | ExBarSeq showed barcodes mostly in soma and sparsely through apical dendrites. Fewer barcodes were seen farther away from the cell body. | -Circular RNA, eg. (Litke and Jaffrey 2019)<br>-AAV variants, Rabies (Chatterjee et al. 2018)<br>-Better targeting motifs<br>-Better RNA stability<br>- Higher RNA expression<br>- Develop transgenic animals with molecular constructs made for multiplex readout<br>-Protein barcodes (Li et al. 2020) |
| Synaptic resolution | Imaging resolution must be able to resolve the location of synaptic sites | Expansion Microscopy with sufficient expansion factor to be able to see synaptic proteins. | - Demonstrate stitching of multiple intact gels together to assemble sections greater than a few cubic millimeters<br>- Automated and roboticized sample handling |
| Molecular interrogation | The technique must be able to stain for multiple endogenous proteins or other cell state markers | This study multiplexes antibody stains against two synaptic markers as well as GFP | - Explore additional antibodies for pre- and post-synaptic proteins including neurotransmitter receptors<br>- Access antibody signal as ExSeq amplicon, eg (Kohman and Church 2020)<br>- Incorporate transcriptional markers (BarSeq) |
| Self contained computational analysis | There must be no minimum training data requirement. Computation should be reliable to non-coders, and readily scale up to whole brain datasets | NMF discovers barcodes in a sample via cell bodies then searches for matching amplicons | - Automate processing directly from the microscope<br>- Create automated instruments |

Our current demonstration of ExBarSeq satisfies three of the four key characteristics. Synaptic staining with a sufficient expansion factor illustrates *synaptic resolution* and *molecular interrogation*. Also, we also developed *self-contained computational analysis* that relies only on the given data. Thus *distance-independent labeling* was the critical characteristic that was not achieved. We also list example future work opportunities for each dimension in **Table 1**.

Moving forward, several approaches could be utilized to improve cell labeling. For near term goals of circuit mapping, vectors could be optimized to increase RNA expression and stability. The use of AAV would enable better targeting of genetically defined cell circuits and the long expression time may produce more uniform subcellular barcode expression. The molecular engineering of barcodes has potential for future innovation; as one example, circular RNA could be pursued which are known to display longer half lives within cells and therefore would be likely to more effectively fill cells. However, it is yet unknown what cellular responses might occur with high expression of exogenous circRNA. Also, improved synapse targeting motifs could be screened to achieve better targeting. For the longer term view toward connectomic goals, new transgenic mice could be created that enable barcode expression throughout all cells in the brain (Li et al. 2020; Marblestone et al. 2014; Kalhor et al. 2018). Lastly, alternative barcoding modalities, such as proteins, could be pursued. The targetability, stability, and long intracellular half lives of proteins make them ideally suited to use as cellular barcodes; however, the ability to write and read high diversity protein libraries is currently limited and needs innovation.

We hope that an enriched era of brain mapping will start when there is an established, integrated set of tools that anyone in the neuroscience community could utilize for their specific scientific question. As transgenic animal models have revolutionized the study of disease (Jaenisch 1975) and, recently, animal-wide development (Kalhor et al. 2018; Tabansky et al. 2013; Veling et al. 2019), so too could engineered animal models be a common platform for brain mapping when paired with a technology such as ExBarSeq and combined with an automated sample handling pipeline. Such an extensible, common technology platform could allow neuroscientists to focus their maximal creative energy on the mysteries of the brain that they seek to understand.

## Acknowledgements

We would like to thank the following people for their contributions over the course of the project: Liam Paninski, Alexei Koulakov, Tai Sing Lee, Sandy Kuhlman, Samuel Inverso, Brian Turczyk, Allison Martin, and Steven Perrault. We acknowledge Microscopy Resources on the North Quad (MicRoN) core at Harvard Medical School for additional access to microscopy equipment. For funding, A.H.M., A.M.Z., E.S.B., G.M.C., and R.E.K. acknowledge IARPA MICrONS (D16PC0008). S.A. was supported by the Howard Hughes Medical Institute (HHMI) fellowship of the LSRF; D.R.G. was supported by an NSF GRFP fellowship; A.T.W. was supported by the Hertz Foundation Fellowship and the Siebel Scholarship; and A.S. was supported by the NIH Neuroimaging Training Program T32 grant 5T32EB001680. E.S.B. was supported by NIH 1R01EB024261, Open Philanthropy, Lisa Yang, John Doerr, NIH 1R01MH123403, NIH R01MH124606, HHMI, Schmidt Futures, and NIH 1U19MH114821.

## Author Contributions

A.H.M., A.M.Z., E.S.B., and G.M.C. initiated the project. D.L., G.L.H., A.C., and R.E.K. performed experiments. D.R.G., R.Z., A.G.X., Y.B., and A.K. performed computational analysis. S.A., G.L.H., X.C., A.T.W., and A.S. contributed to protocol optimization. D.R.G., A.V., R.Z., and R.E.K. contributed to figures. D.R.G., A.V., and R.E.K. wrote the paper. D.R.G. supervised computational analysis. A.V., A.M.Z., E.S.B., G.M.C., and R.E.K. supervised the project. R.E.K. supervised the consortium.

