## Supplemental Information for "Expansion Sequencing of RNA Barcoded Neurons in the Mammalian Brain: Progress and Implications for Molecularly Annotated Connectomics"

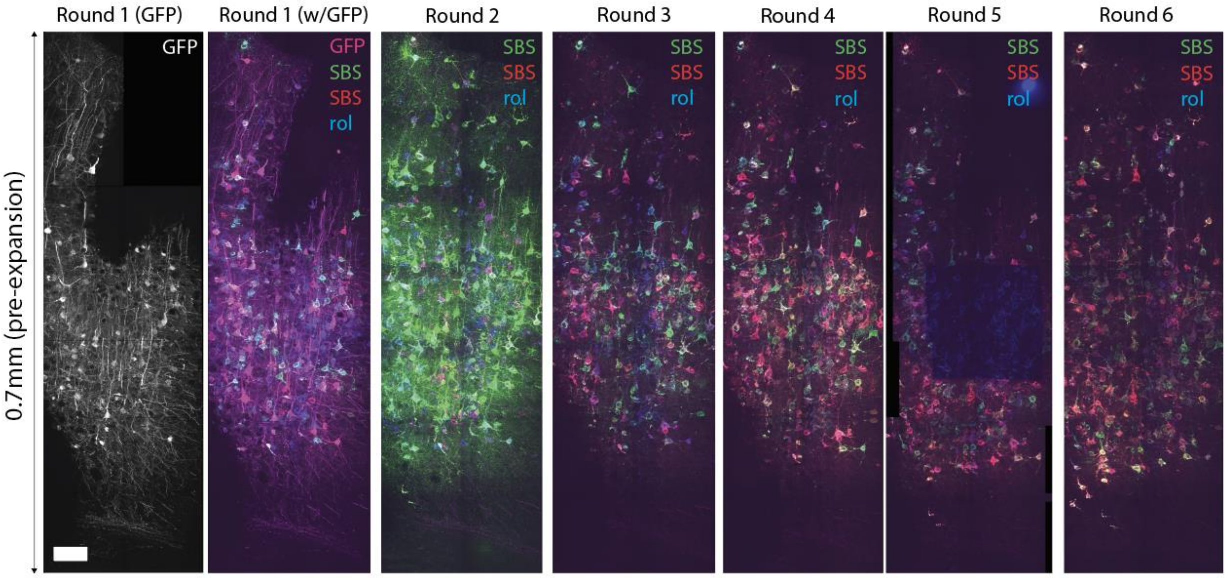


**Supplemental Figure 1 | Overview of six rounds of ExBarSeq data.** All fourteen fields of view are shown across six sequencing rounds. Just the GFP is shown on the far left for clarity on the morphological information at the injection bolus, and then shown combined with the sequencing by synthesis (SBS) color channels. Round 5 had a microscope error which resulted in a fluorophore bleaching in the middle of the sample, rendering Round 5 unusable. In addition to sequencing channels, all RNA barcodes (rol) are shown. Scale bar: 100 µm

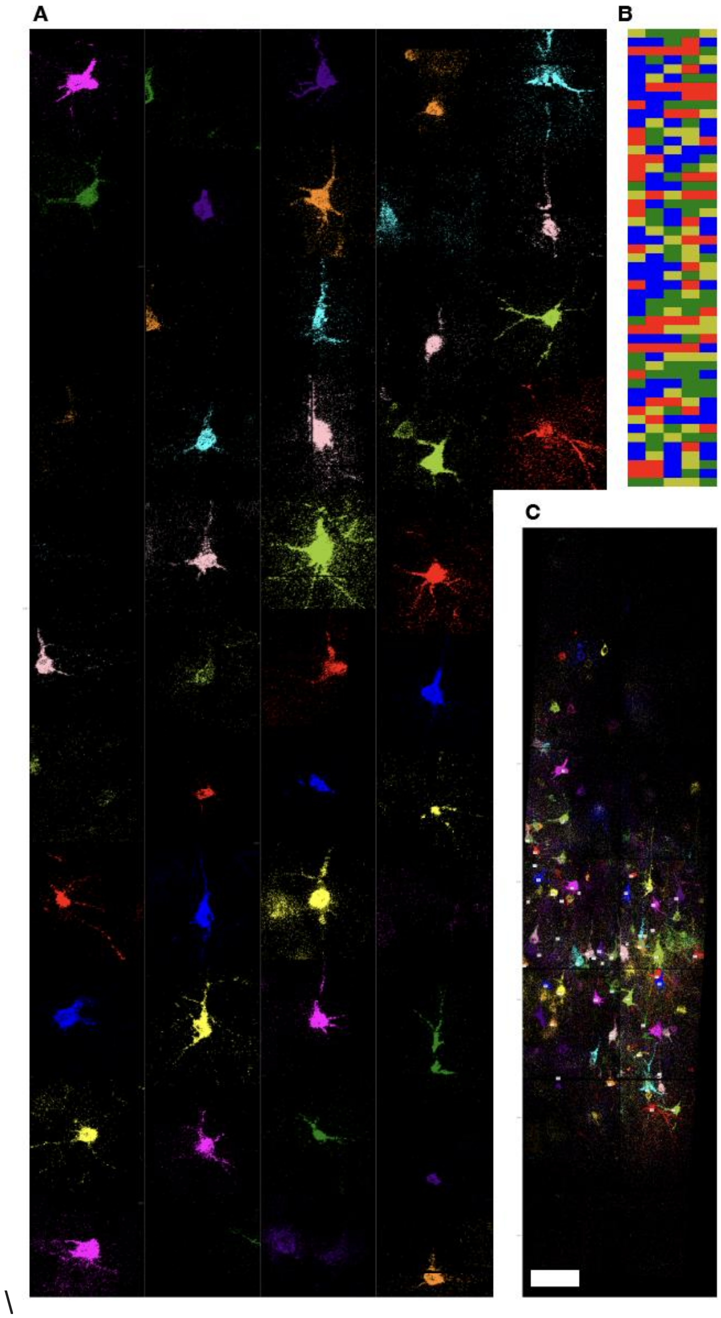


**Supplemental Figure 2 | ExBarSeq computational pipeline automatically extracts neurons**. (A) 49 extracted neurons are shown, illustrating the variety of cellular expression patterns of the MAPSeq virus as determined by the ExBarSeq extraction pipeline. (B) All the 5-base barcodes for the neurons shown in (A) are visualized with the color code in **Fig 3E** showing a diverse set of barcodes. (C) Zoomed out of all extracted neurons discovered. Scale bar: 100 µm

### Methods

#### Solution Recipes

##### Blocking Buffer:

Pierce Protein-Free Blocking Buffer (PFBB) + 0.1% TritonX

##### Crowding Buffer:

| | |
|---|---|
| 500 uL | 20x SSC |
| 500 mg | 40kDa Dextran |
| 5 uL | Triton X-100 |
| 3995 uL | Pierce Protein Free Blocking Buffer |
| **5 g** | **Total** |

##### Reverse Transcription (RT) solution:

| | |
|---|---|
| 20 µL | LNA primer |
| 40 µL | SSIV buffer (5x) |
| 4 µL | dNTP (25mM) |
| 2 µL | BSA (20ug/uL) |
| 5 µL | Ribolock (40U/uL) |
| 10 µL | DTT (0.1M) |
| 10 µL | SSIV (200U/uL) |
| 109 µL | Ultrapure Water |
| **200 µL** | **Total** |

**Hybridization Buffer:**

| | |
|---|---|
| 20 µL | 20x SSC |
| 20 µL | Formamide |
| 160 µL | Ultrapure Water |
| **200 µL** | **Total** |

**Gelling solution:**

| | |
|---|---|
| 188 µL | Monomer solution |
| 4 µL | 4-hydroxy-TEMPO (0.5%) |
| 4 µL | TEMED (10%) |
| 4 µL | ammonium persulfate (10%) |
| **200 µL** | **Total** |

**Monomer solution:**

| | |
|---|---|
| 2.25 mL | Sodium acrylate (380 mg/mL) |
| 0.5 mL | Acrylamide (500 mg/mL) |
| 0.75 mL | N,N′-Methylenebisacrylamide (20 mg/mL) |
| 4 mL | Sodium Chloride (292 mg/mL) |
| 1 mL | PBS (10x) |
| 0.9 mL | Ultrapure Water |
| **9.4 mL** | **Total** |

**Digestion solution:**

| | |
|---|---|
| 150 $\mu$L | 1 M Tris pH 8.0 |
| 6 $\mu$L | 500 mM EDTA, |
| 15 $\mu$L | Triton X-100 |
| 300 $\mu$L | 8 M guanidine HCl |
| 30 $\mu$L | Proteinase K (8 units/mL) |
| 2499 $\mu$L | Ultrapure Water |
| **3 mL** | **Total** |

**Re-embedding solution:**

| | |
|---|---|
| 150 uL | 19:1 Acrylamide: N,N-Methylenebisacrylamide (40%) |
| 10 uL | Tris base (1M) |
| 15 uL | TEMED (10%) |
| 15 uL | APS (10%) |
| 1610 uL | Ultrapure Water |
| **1800 uL** | **Total** |

**Passivation solution 1:**

| | |
|---|---|
| 100 µL | Ethanolamine HCl (4M) |
| 100 µL | MES buffer pH 6.5 (200 mM) |
| 6 mg | EDC |
| 3 mg | NHS |
| **200 µL** | **Total** |

**Passivation solution 2:**

| | |
|---|---|
| 100 µL | Ethanolamine HCl (4M) |
| 100 µL | Sodium borate buffer pH 8.5 (125 mM) |
| **200 µL** | **Total** |

**Padlock solution:**

| | |
|---|---|
| 20 µL | Ampligase buffer (10x) |
| 0.2 µL | /5p/padlock (100 uM) |
| 1 µL | Ampligase (100U/ul) |
| 0.4 µL | dNTP (25mM) |
| 16 µL | RNase H (5U/ul) |
| 20 µL | Phusion DNA polymerase (2U/ul) |
| 5 µL | RiboLock RNase Inhibitor (40U/ul) |
| 20 µL | KCl stock solution (0.5M) |
| 8 µL | formamide |
| 109.4 µL | Ultrapure Water |
| **200 µL** | **Total** |

**RCA solution:**

| | |
|---|---|
| 20 µL | phi29 polymerase (10U/ul) |
| 20 µL | phi29 polymerase buffer (10x) |
| 2 µL | dNTPs (25 mM) |
| 2 µL | BSA (20 ug/ul) |
| 20 µL | glycerol (50%) |
| 1 µL | aadUTP (4mM) |

| | |
|---|---|
| 135 µL | Ultrapure Water |
| **200 uL** | **Total** |

#### Reagents:

#### DNA

| *Name* | *Vendor* | *Sequence* | *Final Concentration* | *Notes* |
|---|---|---|---|---|
| RT primer | IDT | /5ACryd/T+GG+TC+GT+AC+CTCACGACG | 40 uM | Initiates reverse transcription of RNA barcodes |
| Padlock probe | IDT | /5Phos/GTACTGCGGCCGCTACCTAATCCTCTATGATTACTGACTGCGTCTATTTAGTGGAGCCATTGCTATCTTCTTACACGACGCTCTTCCGATCT | 100 nM | Used to padlock barcode cDNA |
| Sequencing Primer | IDT | /5Alex405N/GGAGCCATTGCTATCTTCTTACACGACGCTCTTCCGATCT | 10 nM | Used to simultaneously image and prime amplicons for sequencing |
| Anti A1 azido | IDT | /5ACryd/CCGAATACAAAGCATCAACGAACCGAATACAAAGCATCAACG/3AzideN/ | NA | Conjugate to Bassoon (anti mouse secondary) antibody |

| | | | | |
|---|---|---|---|---|
| Anti A2 azido | IDT | /5ACryd/GGTGACAGGGATCACAATCTAAGGTGACAGGGATCACAATCT/3AzideN/ | NA | Conjugate to Homer1 (anti rabbit secondary) antibody |
| Anti B1 azido | IDT | /5ACryd/TACGCCCTAAGAATCCGAACAATACGCCCTAAGAATCCGAAC/3AzideN/ | NA | Conjugate to GFP (anti chicken) antibody |
| A1 atto550 | IDT | CGTTGATGCTTTGTATTCGG/iSp9//3ATTO550N/ | 100 nM | For imaging Bassoon |
| A2 atto647N | IDT | AGATTGTGATCCCTGTCACC/3ATTO647NN/ | 100 nM | For imaging Homer1 |
| B1 AF488 | IDT | GTTCGGATTCTTAGGGCGTA/3AlexF488N/ | 100 nM | For imaging GFP |

**Antibodies**

| *Name* | *Vendor* | *Catalog number* | *Final Concentration* |
|---|---|---|---|
| Chicken anti-GFP | Millipore | AB16901 | 1:100 |
| Mouse anti-Bassoon | Abcam | ab82958 | 1:200 |
| Rabbit anti-Homer1 | Synaptic Systems | 160003 | 1:200 |
| Donkey anti-Chicken | Jackson ImmunoResearch | 703-005-155 | 1:100 after conjugation |
| Donkey anti-Mouse | Jackson ImmunoResearch | 715-005-151 | 1:100 after conjugation |
| Donkey anti-Rabbit | Jackson ImmunoResearch | 711-005-152 | 1:100 after conjugation |

### Sample Preparation

Unless otherwise stated, all reaction steps were performed with tissue/gels free floating in 200 uL of solution in wells in 48 well plates. PCR tape was placed above each well to ensure no evaporation was occurring.

##### Tissue barcoding and preparation

Viruses preparation and injection were performed according to the MAPseq protocol (Kebschull et al. 2016). For tissue preparation, mice were transcardially perfused with PBS and then 4% paraformaldehyde in PBS. Brains were removed and soaked an additional 24 hours in fixative at 4C. They were then submerged in 30% sucrose at 4C for at least 24 hours in which they were observed to sink to the bottom of the tube. They were then embedded by freezing in OCT using a dry ice/2-methylbutane bath. Samples were sliced on a cryostat at 30 $\mu$m and stored in 70% ethanol until use.

**Reverse Transcription, Staining. and Expansion**

**Reverse Transcription (pre-expansion)**

Tissues were washed several times with PBST to remove the ethanol and then incubated with reverse transcription (RT) solution overnight at 37C.

**Staining protocol (pre-expansion)**

1. Wash slices with PBS and then block in Blocking Buffer for 1 hour.
2. Add primary antibody solution (1:200 Rb Homer, 1:200 Ms Bassoon, 1:100 Chk GFP diluted in Blocking Buffer) and incubate overnight at 4C.
3. Wash with Blocking Buffer 2x for 5 minutes each
4. Incubate in secondary oligo-conjugated antibody solution (1:100 in Crowding Buffer) overnight at room temp
5. Wash with PBS 2x for 5 minutes each
6. Wash with Hybridization Buffer 2x for 5 minutes each
7. Add fluorescent hybridization probes in Hybridization Buffer (0.1µL hybridization probe per 100uL solution) and incubated for 20 minutes at room temperature
8. Wash with Hybridization Buffer 3x for 5 minutes each

**ExM protocol**

The following steps are very similar to ExM1.0 (F. Chen, Tillberg, and Boyden 2015) and explained in detail in (Asano et al. 2018)

1. Place scotch tape on the sides of the slide flanking the tissue

2. Put gelling solution onto the sides of the slice and place a cover slip on top so that the tissue is in between the slide and coverslip

3. Add fresh Gelling solution and incubate for 90min at 37C

a. Remove coverslip and trim excess gel with a razor blade

4. Add gel to Digestion solution and incubate at 37C overnight

**Re-embedding protocol**

The following steps are very similar to (Alon et al. 2021)

1. Add water until gel is fully expanded

2. Add Re-embedding solution, place coverslip in gel, and incubate at 37C for 90 min

3. Trim gel with razor to desired size

4. Add Passivation solution 1 and incubate at room temperature for 2 hours

5. Add Passivation solution 2 and incubate at room temperature for 40 min

6. PBS washes x3

**Probe hybridization and amplification**

1. Add Padlock solution and incubate for 30 min at 37C and then for 45 min at 45C

2. PBST washes for 5 minutes x2

3. Add RCA solution and incubate overnight at room temperature

4. PBST

5. 40ul BS(PEG)9 in 160 µl PBST at RT for 1hr

6. 1MTris-HCl 8.0 wash

7. 1M Tris-HCl 8.0 for 30min at room temperature

8. 2xSSC, 10% formamide x3

For rolony detection – add hybridization probe at 2.5 µM in Hybridization Buffer

For sequencing – add primer at 2.5 µM in Hybridization Buffer, wash, and proceed to sequencing. Refer to (Alon et al. 2021) for more details.

**Sequencing**

Reagents for sequencing were utilized from Illumina NextSeq 500 Mid Output Kits V2 (FC-404-2003) cartridges. We refer to the contents of the largest buffer reservoir as Incorporation Buffer, the purple colored solution as Incorporation Mix, and the solution used to remove fluorescence as Cleave Solution. All reactions were performed in the free floating state using 200 µL of solution per sample. Slices were washed three times with Incorporation Buffer. Then a solution of 50% Incorporation Mix in Incorporation Buffer was added to the slices and incubated at 50ºC for 15 minutes. This step was performed twice each time with fresh solution. Post-incorporation, the slices were washed with Incorporation Buffer at 50ºC for 15 minutes 3 times before imaging. After imaging, slices were incubated with Cleave Solution for 20 minutes at 50ºC. Samples were washed with Incorporation Buffer at 50ºC for 15 minutes 3 times before proceeding to the next round of sequencing.

**Microscopy**

Confocal imaging was performed using a Nikon Ti inverted microscope containing a Yokogawa CSU-W1 spinning disk confocal head with an ORCA-Flash4.0 V3 Digital CMOS camera. Confocal imaging used a 20X water immersion CFI Apo LWD Lambda S objective lens (MRD77200, 0.95 NA) and 405, 488, 561, and 640 nm laser lines.

**Antibody-DNA Conjugation**

To a solution of 100 µL (0.813 nmol) antibody (donkey anti mouse, Jackson ImmunoResearch, 715-005-151) was added 2.00 µL (20.0 nmol) of a 10 mM solution of DBCO-PEG13-NHS in DMSO (Aldrich). After 2 hours, the reaction was purified with a 50k Amicon spin filter (3 spins with PBS, 5 min & 14k x g per spin). The degree of functionalization was calculated by dividing the concentration of dibenzylcyclooctyne (DBCO) by the concentration of the antibody. Concentrations were calculated using Beer's law with absorbances measured using UV-vis spectroscopy (Nanodrop).

Extinction coefficients of 204,000 M-1 cm-1 (at 280 nm) and 12,000 M-1 cm-1 (at 309 nm) were used for DBCO and antibody respectively. 15 µL of the antibody solution (0.359 nmol) was then reacted overnight with 32 µL (10 uM TE, 3.2 nmol) azido oligo used (100 µM). The final product was obtained at a similar concentration to the starting antibody and used without any further purification. This protocol gave comparable results using antibodies against rabbit and chicken (Jackson ImmunoResearch).

### Computational Processing

#### Image Processing

##### Stitching

TeraStitcher (Bria and Iannello 2012) is used for each sequencing round to align all the fields of view (FOVs) to their neighbors. Some pre-processing of the FOVs are needed and explained in detail in the TeraStitcher instructions (FOVs are referred to as “tiles” in TeraStitcher documentation). Briefly, FOVs of images taken from the microscopy are renamed and organized based on its physical position (with the unit of microns). Each FOVs have the shape of 2048 x 2048 pixels with the voxel size of 0.34 µm x 0.34 µm x 0.3 µm. The overlap between tiles is 10%.

The stitching parameters are calculated in TeraStitcher based only on the rolony channel and then later applied on other channels. TeraStitcher outputs the full resolution images and images are then downsampled by 8 fold.

The downsampled images are fed into the ExSeq pipeline to calculate the affine transformation matrix to estimate the gross rotations between imaging rounds in which the same was free-floating. The affine transformation matrix includes the translation and linear warp (rotation and shearing), which can then be used to re-section the original, full-resolution FOVs into “virtual” Fields of View (vFOVs). The advantage of vFOVs is that they have been approximately aligned, meaning that re-calculating the linear warp for a local region will have minimal data loss by maximal overlap of the vFOVs. Furthermore, these vFOVs are beneficial for maintaining standard compute requirements (e.g, 256 GB RAM) for full-resolution analysis. Each vFOV is processed by the ExSeqProcessing pipeline available on Github and described in further detail in (Alon et al. 2021).

#### Barcode Extraction

First, the registered vFOV's two channels sequencing data are quantile normalized and background signal is subtracted. The remaining computation is in two steps: cell body discovery, barcode basecalling, and amplicon alignment.

To locate cell bodies without prior knowledge, we identify bright, highly correlated patches of pixels that have high channel purity (i.e. high signal in one channel relative to others, per round). The data is first partitioned into smaller chunks. Then, we first assign a purity score to each pixel, and then iteratively find regions with the highest cross-correlation to a gaussian with sigma as the rough size of expected cell bodies, similar to (Pnevmatikakis et al. 2016). For each local maxima, we calculate a local rank-1 NMF approximation of the region, and then update the initialization function by removing the factored out signal. This process continues until remaining signal in the region has decreased beyond a threshold percent of the original signal, or a pre-set cell # threshold is reached. After merging across partitions,and removing duplicate candidate cell bodies, the spatial footprint of the candidates is optimized across the full image stack, fixing the channel-across-rounds factored signal. The resulting optimized components are filtered by size, pixel density, and specificity of barcode signal in the spatial footprint area to generate a list of cell body locations and channel-by-round signal vectors.

Barcode basecalling is done with a simple discriminative loss function (De Brabandere, Neven, and Van Gool 2017) by optimizing a set of basecalling weights to map pixels from the same cell body to similar barcodes while mapping pixels from different cell bodies to distant barcodes. By setting a set of initialization weights, we ensured that the weights followed the traditional decoding of the two sequencing channels. The output of this step is an ACTG barcode sequence for each cell body.

Finally, we use the amplicon extraction function of ExSeqProcessing library (Alon et al. 2021) and compare the sequence of the average pixel values per rolony against the dictionary of known barcodes. Perfect match rolonies to known barcodes are then assigned to the respective cell bodies.

Python notebook scripts for the stitching and barcode extraction can be accessed at https://github.com/dgoodwin208/exbarseq
